# Fitness landscape of a dynamic RNA structure

**DOI:** 10.1101/2020.06.06.130575

**Authors:** Valerie WC Soo, Jacob B Swadling, Andre J Faure, Tobias Warnecke

## Abstract

RNA structures are dynamic. As a consequence, mutational effects can be hard to rationalize with reference to a single static native structure. We reasoned that deep mutational scanning experiments, which couple molecular function to fitness, should capture mutational effects across multiple conformational states simultaneously. Here, we provide a proof-of-principle that this is indeed the case, using the self-splicing group I intron from *Tetrahymena thermophila* as a model system. We comprehensively mutagenized two 4-bp segments of the intron that come together to form the P1 extension (P1ex) helix at the 5’ splice site and, following cleavage at the 5’ splice site, dissociate to allow formation of an alternative helix (P10) at the 3’ splice site. Using an *in vivo* reporter system that couples splicing activity to fitness in *E. coli*, we demonstrate that fitness is driven jointly by constraints on P1ex and P10 formation and that patterns of epistasis can be used to infer the presence of intramolecular pleiotropy. Importantly, using a machine learning approach that allows quantification of mutational effects in a genotype-specific manner, we show that the fitness landscape can be deconvoluted to implicate P1ex or P10 as the effective genetic background in which molecular fitness is compromised or enhanced. Our results highlight deep mutational scanning as a tool to study transient but important conformational states, with the capacity to provide critical insights into the evolution and evolvability of RNAs as dynamic ensembles. Our findings also suggest that, in the future, deep mutational scanning approaches might help us to reverse-engineer dynamic interactions and critical non-native states from a single fitness landscape.

## INTRODUCTION

Many RNAs need to fold into defined structures to function. This includes key RNAs in information processing (e.g. rRNAs, tRNAs), RNAs with catalytic activity (ribozymes), and many smaller RNAs (e.g. microRNAs) whose biogenesis depends on base-pairing of a precursor molecule. The need to fold into specific structures and avoid erroneous intra- and intermolecular interactions constrains RNA evolution and evolvability (Chen *et al.* 1999; Umu *et al.* 2016), because at least some mutations will compromise folding, function, and fitness.

Over the last decade, mutational effects on molecular fitness have been elucidated at scale for a handful of model RNAs using deep mutational scanning (DMS) experiments, both *in vitro* (Pitt and Ferré-D’Amaré 2010; Hayden *et al.* 2011; Petrie and Joyce 2014; Kobori and Yokobayashi 2016; Pressman *et al.* 2019; Andreasson *et al.* 2020) and *in vivo* (Zhang *et al.* 2009; Guy *et al.* 2014; Li *et al.* 2016; Puchta *et al.* 2016; Domingo *et al.* 2018; Li and Zhang 2018). These studies have revealed complex fitness landscapes, in which both pairwise and higher-order epistasis are prevalent (Weinreich *et al.* 2013; Lalić and Elena 2015; Bendixsen *et al.* 2017; Domingo *et al.* 2018).

In some instances, mutational effects on fitness and the origins of epistasis can be rationalized with reference to a known (native) structure. It is easy to see, for example, how base-pairing in a conserved helix of a tRNA can be disrupted by a first mutation but then restored by a second, leading to positive epistasis (Li *et al.* 2016). Frequently, however, the molecular foundations of variable constraint and epistasis remain obscure.

Part of the explanation for this likely rests in the fact that RNA structures are dynamic (Ganser *et al.* 2019). As an RNA interacts with itself and its binding partners – during biogenesis, folding, and normal function – conformational changes alter the effective genetic context of a given mutation, i.e. the context that determines mutational impact at a particular point in the life cycle of the RNA. As a consequence, a single static structure, taken as the sole representative from a dynamic conformational ensemble, can only ever act as a partial guide and will sometimes fail to inform on the context(s) in which a particular mutation exerts its effect(s).

DMS experiments allow simultaneous measurement of mutational effects across multiple conformational states, however transient, as long as these states affect fitness (as measured by the experiment). The challenge is to allocate observed patterns of constraints and epistasis to these transient conformational states, which, even if critical for function, are usually unknown and can typically not be extrapolated from knowledge of the native structure.

Here, we investigate the fitness landscape of a dynamic RNA structure that, in our assay, assumes multiple conformational states with known relevance to fitness. We consider a derivative of the group I intron from *Tetrahymena thermophila* (Figure 1A), a self-splicing ribozyme whose functional elements and key catalytic steps have been dissected in great detail using a combination of genetic, biochemical and structural approaches (Cech 1990). To measure molecular fitness and characterize epistatic interactions, we use a previously developed heterologous reporter system where the intron is embedded in a kanamycin nucleotidyltransferase (*knt)* gene (Figure 1B), placed on a plasmid and transformed into *E. coli*. This system couples self-splicing activity to fitness (Figure 1C) as intron removal is required for the reconstitution of the *knt* open reading frame whose translation enables growth in the presence of kanamycin (Guo and Cech 2002).

**Figure 1.**
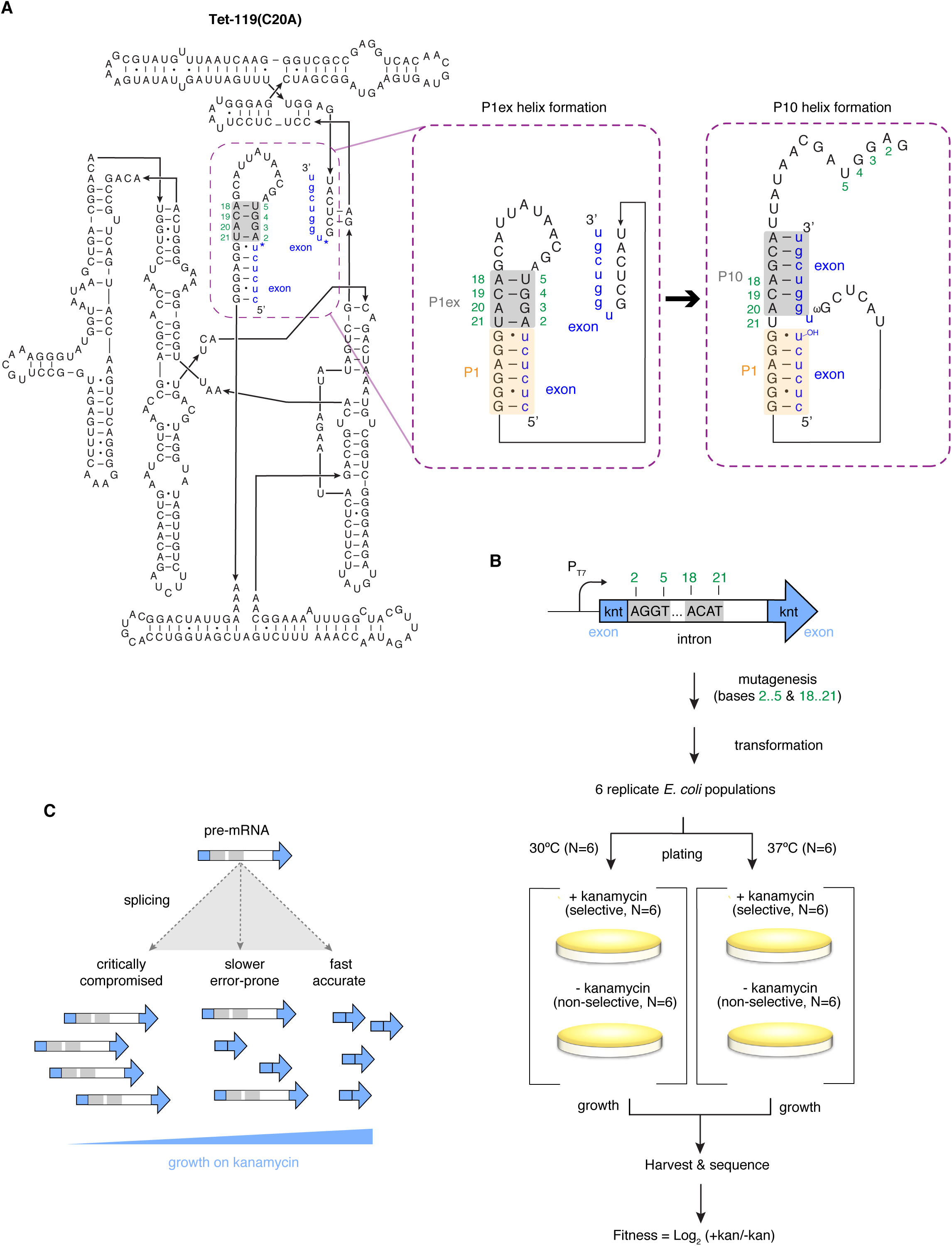
Determining the fitness landscape of a dynamic RNA structure. (**A**) The sequence and secondary structure of the Tet-119(C20A) group I intron with its 5’ and 3’ exonic context. Secondary structure conformations during sequential formation of P1ex and P10 are highlighted in the blow-ups. The two sub-regions that were subjected to mutagenesis (N_2_..N_5_ and N_18_..N_21_) are shaded grey. (**B**) Schematic representation of the *knt*-intron construct, library generation, and selection protocol. (**C**) In the presence of kanamycin, self-splicing activity (molecular fitness) of the group I is coupled to organismal fitness as intron removal is required for reconstitution of the *knt* open reading frame.

We investigate two sub-regions in the intron, N_2_..N_5_ and N_18_..N_21_, which come together to form the P1 extension (P1ex), a 4-bp helix adjacent to the 5’ splice site (Figure 1A). Importantly, following cleavage at the 5’ splice site, P1ex needs to dissociate to allow formation of a second helix (P10), where one half of P1ex (N_18_..N_21_) pairs with bases at the 5’ end of the 3’ exon (Michel et al 1989) (Figure 1A). Constraints on the two sub-regions are therefore asymmetric (with additional constraint on N_18_..N_21_) and pleiotropic (as N_18_..N_21_ function as part of P1ex and subsequently P10). Although the presence of neither P1ex nor P10 is strictly required for splicing (Been and Cech 1985; Price and Cech 1988; Cech 1990), both helices contribute to splicing efficiency, as they facilitate splice site alignment and exon ligation and reduce non-productive alternative interactions, including the use of cryptic splice sites (Michel *et al.* 1989; Suh and Waring 1990; Narlikar *et al.* 2000; Bell *et al.* 2004; Karbstein *et al.* 2007). Mutations in P1ex and P10 have previously been shown to affect rates of catalysis at different stages of splicing (Doudna *et al.* 1989; Guo and Cech 2002; Bell *et al.* 2004; Karbstein *et al.* 2007), which is relevant for *knt* production and, subsequently, fitness (Guo and Cech 2002). Prior work has also provided *prima facie* evidence for antagonistic pleiotropy, inferring from a small collection of individual mutants that overly stable pairing in P1ex might be selected against because it impedes dissociation and therefore P10 formation (Guo and Cech 2002; Michael A Bell *et al.* 2004).

Measuring fitness for a large number of intron genotypes that vary at N_2_..N_5_ and N_18_..N_21_, we dissect the resulting fitness landscape to demonstrate that fitness effects of specific mutations can be allocated to distinct conformational states and that DMS data can be used to investigate pleiotropic trade-offs at sites involved in more than one fitness-relevant structure. Our results provide a proof-of-principle that DMS simultaneously captures fitness effects arising from multiple transient conformational states. They also suggest that, in the future, DMS could be used alongside evolutionary analysis, structural modelling, and biochemical approaches to infer transient states at scale.

## RESULTS & DISCUSSION

We used targeted saturation mutagenesis via overlap extension PCR to generate a large library of intron variants (see Methods), using a previously characterized mutant with high splicing activity [Tet-119(C20A), Figure 1A] as our master sequence. Introns differ in the two sub-regions N_2_..N_5_ and N_18_..N_21_ but are otherwise isogenic. The library was transformed into *E. coli* and each biological replicate split into four aliquots, which were spread on agar plates that did or did not contain kanamycin and incubated at either 30°C and 37°C (Figure 1B, Methods). After overnight incubation, genotype frequencies under selective and non-selective conditions were assayed via high-throughput amplicon sequencing (see Methods).

Under non-selective conditions (without kanamycin, *-kan*), where production of functional KNT protein is not required for survival, our library is nearly combinatorially complete. Across 6 biological replicates and 31,269,777 sequencing reads (at 30°C, Table S1), we fail to detect only 3 of all 4^8^=65,536 possible genotypes (>99.99% completeness). As a consequence of the library generation protocol, and similar to prior work (Pitt and Ferré-D’Amaré 2010), sequences closer to the starting template are more common, increasing our power to investigate sequence space closer to the splice-competent master genotype (Figure S1, Methods).

Different genotypes with higher or lower fitness can be thought of as conceptually equivalent to different transcript species that increase or decrease in abundance. We therefore analyzed the data using a method commonly employed for counts-based differential expression analysis (DESeq2) (Love *et al.* 2014). This approach has several advantages, including well-established statistical foundations to determine significant changes in the face of biological variability and simplicity of implementation. We note that fitness estimates derived using DESeq2 are highly correlated to estimates from an alternative method (Bolognesi *et al.* 2019) that explicitly models the main sources of variability in DMS data (r^2^=0.91, P<2.2*10^−16^; Figure S1, Methods).

Under selective conditions (+*kan*), colony formation is much reduced (Figure S2) and the majority of genotypes (42193/65536=64%) experience a significant drop in frequency (at P_adj_<0.05), while only 6.5% (4286/65536) become significantly more common, leading to a precipitous decline in overall genotype diversity (Figure 2A,B). Changes to the composition of the genotype pool are similar across replicates, as quantified using Bray-Curtis dissimilarity (Figure 2C). Individual P1ex genotypes previously found to exhibit increased splicing efficiency have concordant effects in our assay (Figure S3). Similar to the fitness landscapes of other RNAs and proteins, the distribution of fitness effects across genotypes is bimodal [reviewed in (Kemble *et al.* 2019)] and average fitness decreases as the number of mutations away from the master sequence (=Hamming distance) increases (Figure 2A,D).

**Figure 2.**
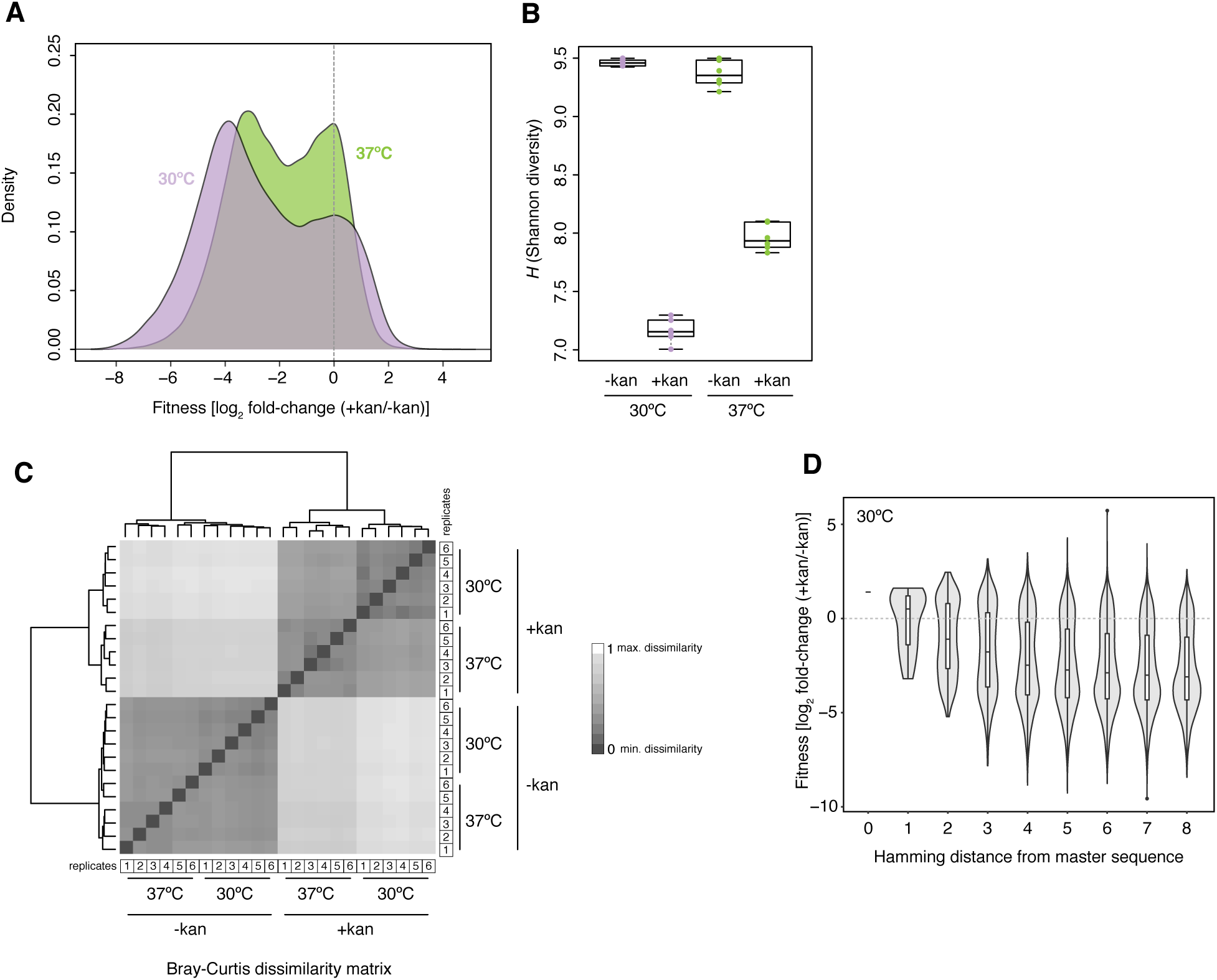
Fitness across intron genotypes. (**A**) Distribution of fitness effects at 30°C and 37°C. (**B**) Shannon diversity of intron genotype pools under different conditions. (**C**) Similarity in genotype pool composition across all replicates and conditions measured as Bray-Curtis (BC) dissimilarity, where BC=1 indicates maximum dissimilarity between samples. (**D**) Fitness of intron genotypes at 30°C as a function of Hamming distance (i.e. the number of mutational steps away from the master sequence).

### Fitness effects across mutant genotypes support selection against excess stability in P1ex

Prior work on both tRNA and snoRNA found fitness defects to be more pronounced at 37°C compared to 30°C (Puchta *et al.* 2016; Li and Zhang 2018), consistent with destabilization of folded structures as a key determinant of mutant fitness. We observe the opposite (Figure 2A). While fitness estimates for individual genotypes are highly correlated between 30°C and 37°C (Figure S4, ρ=0.75, P<2.2*10^−16^), fitness impacts are quantitatively milder, on average, at the higher temperature. This is in line with suggestions that excess stability of P1ex secondary structure compromises efficient splicing (Guo and Cech 2002), as kinetic traps should, on average, be easier to escape and misfolding issues be less severe at 37°C. In support of this explanation, we find greater predicted stability and higher GC content to be associated with larger decreases in fitness (Figure 3A,B; Methods). At the same time, genotypes that cannot form *any* on-target base-pairs also exhibit low fitness (0 strong/weak base-pairs in Figure 3C). In contrast, genotypes where helices *are* formed, but the constituent base-pairs are weak (A-U), as found in the *T. thermophila* native structure (Figure S5), typically do well (Figure 3C).

**Figure 3.**
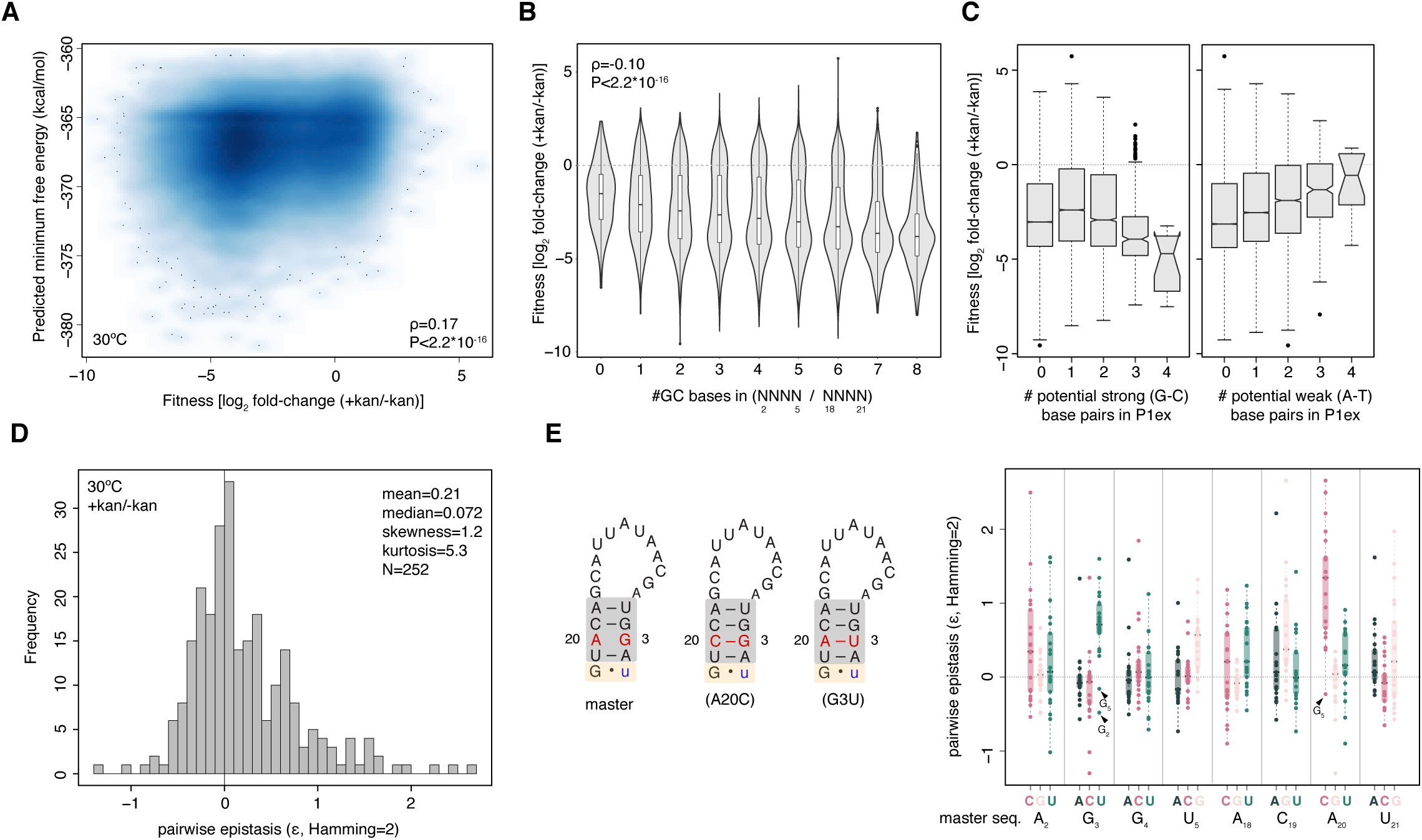
Causes and correlates of variable fitness across intron genotypes (**A**) Fitness weakly correlates with predicted minimum free energy of the intron. Note that the predicted minimum free energy of the master sequence ΔG = -362.8 (B) Fitness varies according to the number of guanosine or cytosines (#GC) in the N_2_..N_5_ and N_18_..N_21_ regions. (**C**) Fitness varies as a function of the number of strong or weak base-pairs that could be formed in P1ex assuming that base-pairing follows the established master/wildtype pattern (see Figure 1A). (**D**) Distribution of pairwise epistasis values for genotypes that are two mutations away from the master sequence (Hamming distance = 2). ε values above 0 indicate positive epistasis, those below 0 indicate negative epistasis. (**E**) Pairwise epistasis for genotypes in (D) by position and mutation. Diagrams on the left highlight the N_3_/N_20_ couple, where mutations that are predicted to lead to base-pairing are associated with positive epistasis.

The need to avoid an overly stable P1ex helix is further evident when looking at patterns of epistasis. In contrast to most other RNA DMS studies (Bendixsen *et al.* 2017), we observe an enrichment for positive rather than negative pairwise epistasis when considering single and double mutations away from the master sequence (Figure 3D). In some instances, positive epistasis corresponds to cases where a base-pair is broken by each of two individual mutations but restored when these mutations are combined. However, we observe multiple cases of strong positive epistasis that do not conform to this model. Notably, many such cases involve A20C and G3U (Figure 3E), the only two mutations capable of generating a helix with four paired bases. Any further mutation elsewhere in the two sub-regions will abolish perfect complementarity in P1ex. Almost always, the reduction in fitness upon adding this second mutation is less severe than expected under an additive model of mutational effects, in line with selection against excess stability. This highlights that positive epistasis can result not only from selection to maintain base pairing but also from selection to prevent it.

### Machine learning facilitates allocation of mutational effects to distinct conformational states

Although simple metrics like stability and GC content are related to fitness, they are overall poorly predictive (GC content: ρ=-0.10; predicted stability: ρ=0.17, Figure 3A,B), suggesting a more complex landscape of constraint than one exclusively defined by a P1ex structural stability threshold. To better understand how specific mutations affect fitness and whether they do so in a P1ex and/or P10 context, we sought to determine the contribution of individual nucleotides to fitness systematically and in a genotype-specific manner. To this end, we trained extreme gradient boosted decision tree (XGboost) models (Chen and Guestrin 2016) to predict fold-changes (+*kan* vs. *-kan*) solely from nucleotide identities at N_2_..N_5_/N_18_..N_21_. For both 30°C and 37°C, we find that fold-changes predicted from the models are well correlated with observations (30°C ρ=[0.63, 0.84], P<2.2×10^−16^; 37°C ρ=[0.63-0.83], P<2.2×10^−16^, see Methods for calculation of correlation ranges). We estimate that these models account for ∼80% of the explainable genetic variance. Providing additional RNA-wide properties as features for prediction (e.g. RNAfold-predicted stability or ensemble diversity) does not improve model performance (Table S2), suggesting that the models capture key emergent properties from the underlying primary sequence. In addition, confining analysis to genotypes whose change in relative abundance was judged significant by differential abundance analysis, does not improve prediction accuracy (Table S2). This suggests that noise from lowly abundant genotypes does not compromise predictive power and even contains latent information that enables more accurate prediction across genotype space.

The contribution of individual features to prediction accuracy can be assessed globally by considering the gain in classification accuracy when a leaf in the tree is split according to that feature. However, computing such *gains* does not provide directionality of effect nor the ability to assess contribution locally, i.e. on a genotype-by-genotype basis. We therefore additionally computed Shapley additive explanation (SHAP) values (Lundberg and Lee 2017; Lundberg *et al.* 2020), which provide a framework for interpreting the impact of individual features on model prediction in a machine learning context, and contain information about both sign and magnitude of the contribution.

In our case, a feature corresponds to having or not having a particular nucleotide, e.g. a cytosine, at a given site, e.g. N_21_. In some instances (e.g. G_5_, Figure 4A), nucleotide identity affects fold-change prediction consistently in the same direction across genotypes, although the precise contribution might vary from genotype to genotype (equivalent to magnitude rather than sign epistasis). In other cases (e.g. U_3_, Figure 4A), the identity of a nucleotide at a particular site only substantively contributes to predictions for a small number of genetic backgrounds.

**Figure 4.**
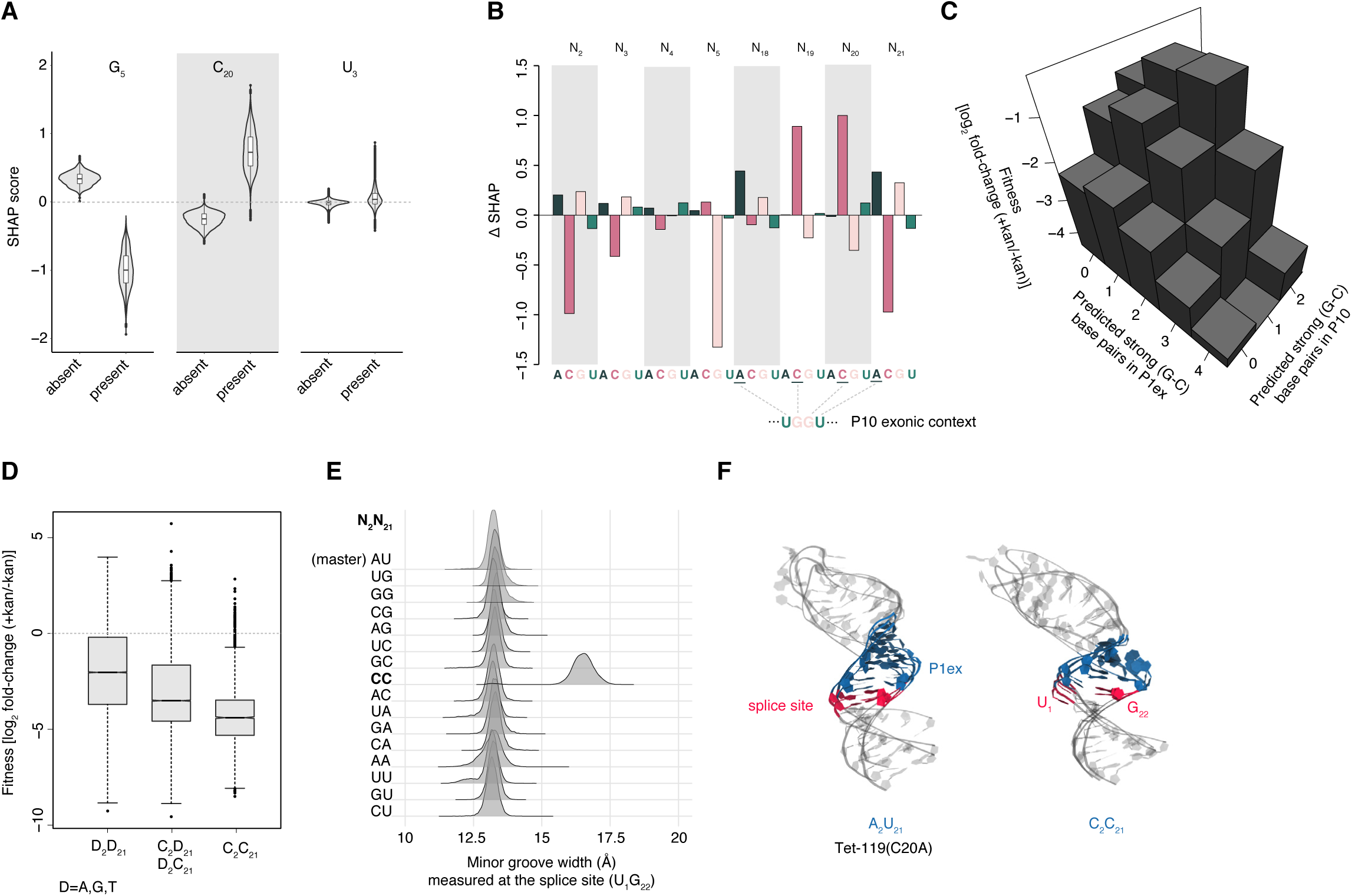
Assessing the contribution of individual nucleotide identities to fitness across multiple structural conformations. (**A**) Contribution to XGBoost-predicted relative fitness across all intron genotypes, as measured by Shapley’s additive explanation (SHAP) scores, of three example site/nucleotide features. More positive SHAP scores are associated with higher fitness. (**B**) The average contribution across all genotypes of all individual site/nucleotide features, measured as ΔSHAP = SHAP_present_ - SHAP_absent_, where SHAP_present_ and SHAP_absent_ correspond to the mean SHAP score of all genotypes where a given nucleotide at a given site is present and absent, respectively. (**C**) Fitness landscape as a function of the number of strong (G-C) base-pairs that can form in P1ex and P10, assuming bases are aligned as they are in the master/wildtype structure (see Figure 1A). (**D**) Fitness as a function of N_2_/N_21_ genotype, with a focus on cytosines. (**E**) Minor groove width associated with different N_2_/N_21_ genotypes as determined using molecular dynamics simulations (see Methods). (**F**) Three overlaid representative conformations of the P1/P1ex helix (randomly sampled from the final 50 ns of each simulation) for the master sequence nd the C_2_/C_21_ genotype.

Figure 4B summarizes the average contribution of each site/nucleotide feature to the prediction by computing ΔSHAP, defined here as the mean SHAP value across genotypes where a given nucleotide at a given position is present minus the mean SHAP value across genotypes where the nucleotide at the same position is absent. Notably, the strongest positive contributions involve nucleotides that allow on-target base-pairing during formation of P10 (A_18_, C_19_, C_20_, A_21_, Figure 4B). This suggests that, even though not essential for splicing (Been and Cech 1985), P10 pairing is a major driver of differential fitness in our system. In contrast, there are no strong positive contributions from nucleotides exclusive involved in P1ex (i.e. N_2_..N_5_). This supports earlier models, which argued that P1ex function is largely independent of sequence as long as minimal structural requirements (including avoidance of excess stability) are satisfied (Doudna *et al.* 1989; Allain and Varani 1995; Karbstein *et al.* 2007). Rather, N_2_..N_5_ is principally characterized by negative constraints, where the presence of specific nucleotides is associated with decreased fitness (Figure 4B).

One such constraint involves bases N_2_ and N_21_, where the presence of cytosines is associated with a strong negative contribution to fitness (Figure 4B, Figure S6). This observation is consistent with prior experiments in the wildtype P1/P1ex context (Figure S5) where an 80% (40%) decline in splicing activity was observed when A_2_-U_21_ was replaced with G_2_-C_21_ (C_2_-G_21_) (Doudna *et al.* 1989). We find fitness defects to be particularly pronounced when cytosines are present at both these sites (C_2_/C_21_, Figure 4D). In the master and wild-type *T. thermophila* sequence, N_2_ and N_21_ form a base-pair directly adjacent to the splice site U_1_-G_22_ (Figure 1A, 3E). We therefore suspected that cytosines at these positions might disturb splice site geometry. To investigate this further, we carried out molecular dynamics simulations (see Methods) of all 16 possible N_2_/N_21_ combinations in an otherwise isogenic Tet-119(C20A) context. Considering a catalogue of features (Lu and Olson 2003) that describe base-pairing geometry (stagger, roll, twist, etc. see Methods) we find that C_2_/C_21_ – uniquely – leads to a radical structural deformation of minor groove geometry (Figure 4E,F; Figure S7; File S1), as the splice site U_1_ rotates out of the helix core and G_22_ mis-pairs with C_2_. This likely disturbs splice site definition and key tertiary contacts between the P1 substrate and the catalytic core of the ribozyme (Strobel and Cech 1993; 1995; 1996; Strobel *et al.* 1998), consistent with poor splicing.

Finally, G_5_ makes a strong negative contribution to fitness, both on average and across genotypes (Figure 4A,B). It is interesting to note in this regard that in many naturally occurring introns, including the native *T. thermophila* intron (Figure S5), no pairing is observed at N_5_-N_18_ resulting in a P1ex helix that is only three bases long. This suggests that having a base-pair at this position and/or extending the helix beyond three bases often interferes with efficient splicing (Figure S6). However, unlike in the case of N_2_-N_21_, the negative contribution of G_5_ is not mirrored on the other side of the helix (at N_18_); we therefore predict that G_5_ might have negative fitness consequences outside the P1ex context that remain to be deciphered.

### Asymmetric fitness effects allow inference of pleiotropy

As described above, we detect mutational effects that are attributable to either P1ex or P10 formation. At the same time it is also evident that trade-offs exist to enable the successive formation of the two structures, as illustrated by the divergent preferences for strong base-pairs (Figure 4C). Given its pleiotropic role in participating in both P1ex and P10, N_18_..N_21_ is at the center of this trade-off and has to satisfy an additional layer of constraint. We asked whether this may be reflected in the relative contributions that different site/nucleotide features in N_2_..N_5_ versus N_18_..N_21_ make to predictions. We find this to be the case: a significantly larger proportion of gains in the model is attributable to N_18_..N_21_ (Figure 5A). This asymmetry is also reflected in patterns of epistasis. When we consider pairwise interactions within N_2_..N_5_ (with N_18_..N_21_ fixed as ACAU), within N_18_..N_21_ (with N_2_..N_5_ fixed as AGGU) or across helices (with one mutation each in N_2_..N_5_ and N_18_..N_21_), we find a tendency for positive epistasis to be more prevalent within N_18_..N_21_ than cross-helix and particularly compared to N_2_..N_5_ (Figure 5B, Wilcoxon text, P<0.1) Thus, positive epistasis is more common, on average, for mutations at nucleotides N_18_..N_21_, consistent with pleiotropic constraint. Distinct landscapes of epistasis in N_2_..N_5_ versus N_18_..N_21_ are also evident when we consider higher-order epistasis by computing the correlation of fitness effects (γ) (Ferretti *et al.* 2016) at different Hamming distances from the master sequence. Finally, to further illustrate asymmetric fitness effects across the P1ex helical divide, we carried out a simple mirror test, where we compare the fitness of a given genotype (e.g. A_2_AAG_5_/C_18_TTT_21_) to its mirror image across the helix axis (here T_2_TTC_5_/G_18_AAA_21_). To provide a fair comparison, we only considered genotypes and their mirror genotypes that are at equal Hamming distance (d=2) from the master sequence. In line with strongly asymmetric fitness effects motifs, we find only a weak, non-significant correlation between the fitness of mirrored genotypes (ρ=0.21, P=0.4; N=19). These results serve as a reminder that, even though restoration (e.g. flipping a G-C to a C-G base-pair) is commonly used to demonstrate the importance of base-pairing and helix formation, two sides of any given helix need not necessarily be equivalent. In fact, we expect asymmetry to be common, caused by differential involvement in folding intermediates and alternative conformational states, but also strand-specific modifications and interactions with chaperones and other proteins and RNAs. Asymmetric effects are likely prevalent even in helices where base-pairing is of pre-eminent concern. tRNAs, for example, are post-transcriptionally modified and interact with proteins (e.g. tRNA synthetase on acceptor) in an asymmetric manner.

**Figure 5.**
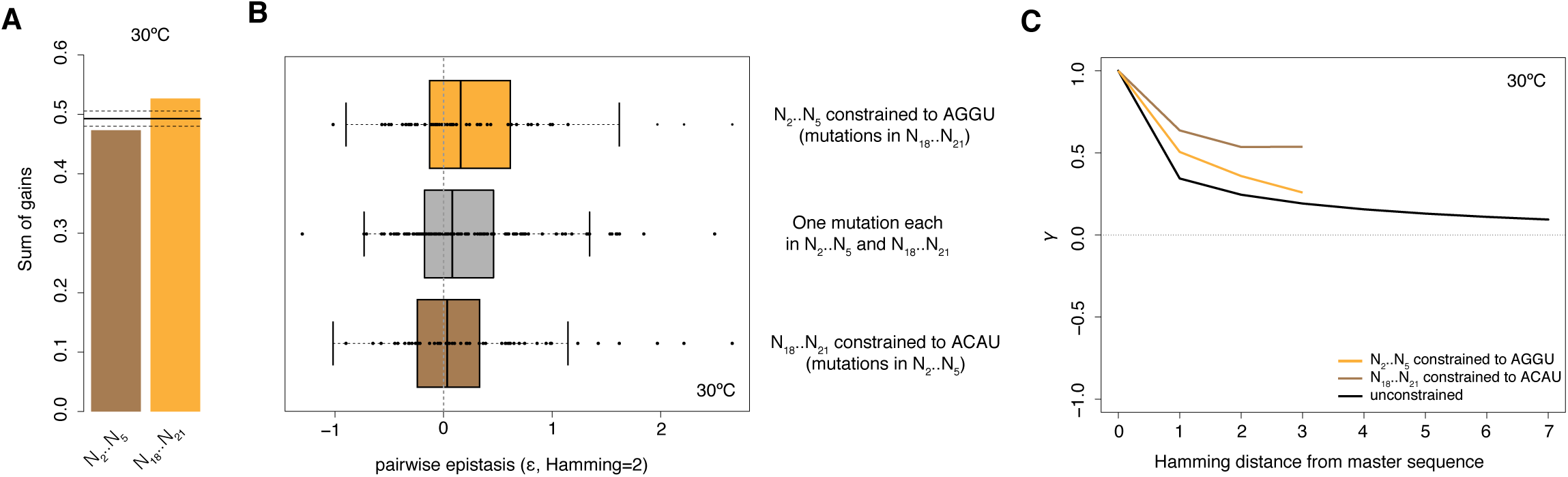
Asymmetric fitness effects across the N_2_..N_5_ and N_18_..N_21_ sub-regions. (**A**) Proportion of gains in the model (see main text) contributed by site/nucleotide identity features at N_2_..N_5_ and N_18_..N_21_. The solid line corresponds to the mean contribution made by a sub-region across 100 random samples, where individual gains are randomly shuffled across site/nucleotide identity features. Dashed lines correspond to 95% confidence intervals. (**B**) Pairwise epistasis for double mutants where both mutations are located in N_18_..N_21_ (orange), both mutations are located in N_2_..N_5_ (brown), or N_2_..N_5_ and N_18_..N_21_ carry one mutation each (grey). (**C**) The correlation of fitness effects (γ) of intron mutants at various mutational distances from the master sequence.

Our study provides a proof-of-principle that DMS experiments can capture multiple fitness-relevant conformational states, including transient states, simultaneously, providing a window onto the fitness of RNAs in their true ensemble state. This capacity to capture multiple structural states in a one-pot experiment brings both opportunities and challenges. Challenges, because mutant fitness need not be interpretable in context of single (native) structure. In fact, mapping fitness effects onto a single native structure might prove misleading at sites where a dominant contribution to fitness comes from non-native or transient conformations or where mutational effects are pleiotropic. At the same time, capturing ensembles brings opportunities: data from DMS experiments might help us identify residues whose contribution to fitness is large but not easily explained when considering the native structure and prioritize these residues for follow-up studies. When used in conjunction with tools to probe and predict RNA structure and function, DMS experiments might, ultimately, even allow us to reverse-engineer dynamic interactions and critical non-native states from a single fitness landscape and provide a better, ensemble-based understanding of RNA evolution and evolvability.

## METHODS

### Construction of mutant intron library

The plasmid backbone of Tet-119 is derived from *E. coli-Thermus thermophilus* shuttle vector pUC19EKF-Tsp3 (Wayne and Xu 1997), which contains a ColE1 *ori*, an ampicillin resistance marker gene, and the *knt-intron* sequence under the control of a *slpA* promoter (Guo and Cech 2002). The *knt-intron* construct was made previously by inserting the intron sequence at nucleotide 119 downstream of the translational start site of *knt*. To maintain base-pairing with the 3’ exon to form P10 and so as not to introduce amino acid substitutions into KNT, nucleotides 15-20 were altered from 5’-TACCTT-3’ (in the wild-type T. *thermophila* intron variant) to 5’-ACGACC-3’. Due to the change in nucleotides 19-20 from 5’-TT-3’ to 5’-CC-3’, nucleotides 3-4 were altered from 5’-AA-3’ to 5’-GG-3’ to maintain base-pairing within the P1ex region. However, *E. coli* strains bearing this intron variant were not viable when challenged with kanamycin, indicative of insufficient splicing activity (Guo and Cech 2002). Tet-119(C20A) was subsequently identified in a screen for mutants that rescued the splicing defect (Guo and Cech 2002).

Upon receipt of Tet-119(C20A), a gift from Feng Guo (UCLA), we amplified the entire *knt-intron* sequence (using primers knt-rz-f and knt-rz-r, Table S3) and subcloned it into the NdeI/XhoI sites of a pET-22b(+) plasmid (Merck Millipore) so that its expression is driven by an IPTG-inducible T7 promoter. To make the mutant library, all eight nucleotides in the two sub-regions were mutated into all possible substitutions (4^8^ variants) using overlap extension PCR (Ho *et al.* 1989; Williams *et al.* 2014) coupled with oligonucleotides containing mixed bases at these sites (Figure S8, Table S3). Note that this procedure, in contrast to protocols employing doped oligonucleotides, will preferentially amplify sequences closer to the starting template as oligos closer to the starting template will bind the template better during PCR.

Oligonucleotides were from Integrated DNA Technologies, and all PCRs were carried out using Q5 High-Fidelity DNA polymerase (New England Biolabs). All DNA fragments were purified from agarose gel (Monarch DNA Gel Extraction kit, New England Biolabs) to reduce carry-over of residual contaminants.

The mutated pool of introns was then ligated into pET-22b(+), and the ligated products were electroporated into competent *E. coli* DH5a (New England Biolabs) cells according to standard procedures (Sambrook and Russell 2012). After electroporation, cells were recovered in SOC medium at 37°C for 1 hour. Recovered cells were then grown on LB agar containing 100 ug/mL carbenicillin at 37°C for 16 hours. The next day, the total number of transformed colonies was estimated to be ∼5.5 × 10^5^, corresponding to at least 8-fold oversampling of the target library size of 4^8^ variants. All transformed colonies were scraped off the agar plates and pooled in 10 mL LB + 100 ug/mL carbenicillin. Half of the pooled cells were archived at -80°C, and the remaining half was harvested for plasmid extraction (QIAprep Spin Miniprep).

### Growth under selective and non-selective conditions

The extracted plasmids from the mutant library were re-electroporated into *E. coli* BL21(DE3) as previously described. For each transformation, 13 fmol of the plasmid library (corresponding to 59 ng) was mixed with 100 uL of electrocompetent bacterial suspension. After electroporation, cells were recovered in SOC medium at 37°C for 1 hour prior to a brief centrifugation (2,500xg, 5 min). The supernatant was removed, and the cells were washed gently with LB. After resuspending the washed cells in 0.5 mL LB, half of the suspended cells (0.25 mL) were used for experiments at 37°C, the other half for experiments at 30°C. For each temperature, a 125-uL aliquot was spread on an LB agar containing 25 ug/mL kanamycin, while another 125-uL aliquot was spread on an LB agar without kanamycin. Other supplements in both media, were 100 ug/mL carbenicillin, 50 uM IPTG and 0.2% rhamnose. Agar plates were then incubated overnight at either 37°C or 30°C. A total of six replicate transformations were carried out, but with only two replicate transformations being conducted in the same day. After incubation, colonies that formed on the agar plates with or without kanamycin were scraped off and pooled using 3 mL LB containing 100 ug/mL carbenicillin accordingly. An 1-mL aliquot of the pooled bacterial suspension was used for plasmid extraction (QIAprep Spin Miniprep) whereas the remaining pooled aliquot was archived at -80°C.

### Library preparation and sequencing

An aliquot (3 fmol each) of the plasmids extracted from the selected and non-selected populations was used for PCR (24 cycles) to amplify a 204-bp sequence spanning the P1ex region using a pair of adapter-linked primers (C20Aseq-f and C20Aseq-r, Table S3). The resulting amplicons from each replicate/strain were cleaned up using the Monarch PCR & DNA Cleanup kit (New England Biolabs). Next, Illumina indices (Nextera XT dual indexing) were incorporated into the adapter-linked amplicons in a second PCR (8 cycles), and the resulting index+adapter-linked amplicons were purified using Ampure XP beads. Index incorporation was confirmed with Agilent Bioanalyser HS-DNA. After quantifying the DNA concentration of the index+adaptor-linked amplicons using Qubit assays analysis (High Sensitivity DNA Assay), each was normalized to 2.5 nM and then combined to make an equimolar pool. The amplicon pools were subjected to 100-bp paired-end sequencing on an Illumina HiSeq 2500 v4 sequencer. To guard against batch effects, we sequenced samples according to a balanced design where each of the 24 samples (6 replicates x 2 temperatures x 2 conditions), along with samples from other conditions not described in this manuscript, was split into three, and one third each allocated to one of three HiSeq lanes for sequencing. Split samples cluster tightly together on PCA, suggesting that batch effects are negligible (not shown). Raw reads have been deposited in the NCBI Sequence Read Archive under accession PRJNA636762. Read/genotype counts after filtering (see below) are provided in Table S1.

### Read processing and fitness estimates

We quality-filtered reads and estimated fitness using two different pipelines. In the first pipeline, we treated the data as one would when conducting a differential expression experiment, where individual genotypes correspond to individual RNA species in a complex pool of transcripts. Reads were trimmed using Trimmomatic v 0.35 (HEADCROP:5 MINLEN:95) and subsequently filtered for base quality >=30 at the mutated bases. Imposing stringent quality cut-offs across the untargeted backbone does not affect results and leads to the removal of many more reads and is needlessly conservative since most deviation here should be owing to sequencing errors. The relative fitness of each genotype (along with adjusted significance values, P_adj_) was then estimated using DESeq2 (implemented in R) as log2-fold change in abundance of a given genotype in six replicates treated with kanamycin compared to six replicates without kanamycin.

For comparison, fitness estimates were computed with DiMSum v0.3.2.9000 (https://github.com/lehner-lab/DiMSum) (Bolognesi *et al.* 2019), which derives final fitness estimates as an error-weighted sum of replicate fitness values, after computing wildtype-normalized fold changes at the replicate level. DiMSum was run with the following parameters: *cutadapt5First*: GGGGATGATGTTAAGGCTATTGGTGTTTATGGCTCTCT, *cutadapt5Second*: CGGTCTTGCCTTTTAAACCGATGCAATCTATTGGTTTAAAGACTAGCTACCAGTGCATGCCTGATAACTTTTCCCTCC, *cutadaptCut3Second*: 1, *cutadaptMinLength*: 20, *cutadaptErrorRate:* 0.2, *usearchMinlen*: 20, *wildtypeSequence*: AGGTagcaatattacgACAT, *maxSubstitutions*: 8. As highlighted above, fitness estimates are highly concordant between the two pipelines (Figure S1). Fitness estimates for all genotypes from both methods are provided in Table S1.

### Computation of summary measures

Shannon diversity and Bray-Curtis dissimilarity were calculated using the *diversity* (index=“Shannon”) and *vegdist* (method=“bray”) functions from the R package *vegan*. Skewness and kurtosis were calculated using the *skewness* and *kurtosis* functions from the R package *moments*. To allow direct comparison to prior results (Bendixsen *et al.* 2017), pairwise epistasis was calculated as log_10_(*f*_*master*_*\*f*_*m1,2*_*-f*_*m1*_*\*f*_*m2*_), where *f*_*master*_ is the fitness of the master sequence and *f*_*m1*_, *f*_*m2*,_ and *f*_*m1,2*_, are the fitness values of the two single-nucleotide mutants and the double mutant, respectively, as calculated by the DiMSum pipeline. Note that fitness in this pipeline is evaluated relative to the master sequence whose fitness is set to 1.

### Computation of RNA structural features

Minimum free energies (MFE) of the different intron genotypes was computed using RNAfold from the Vienna package (v2.4.3, --noPS -p -d2 --MEA -T 37/30), using the intron with ±10 flanking nucleotides, which is sufficient for splicing (Price *et al.* 1987). Results are qualitatively identical when we consider the intron along with the entire *knt* open reading frame instead (not shown).

### Machine learning

Extreme gradient boosted (XGBoost) decision trees were implemented using the *xgboost* and *caret* packages in R, with nucleotide identities encoded via one-hot encoding. Two-thirds of the genotypes were used for training and one third for testing, with 5-fold cross-validation. Hyperparameters were tuned via grid search [nrounds = c(100, 200, 500, 1000), eta = c(0.01,0.05,0.1, 0.3), max_depth = c(4,6,8, 10), subsample = c(0.5, 0.75, 1.0), min_child_weight = c(5, 10, 20)]. Two parameters, colsample_bytree and gamma were set to 1. Models were then using xgbTree using the RMSE metric to minimize (method=“xgbTree”, objective = “reg:linear”, metric=“RMSE”). Predictions are based on the best parameters after tuning. We also carried out equivalent training for subsets of the data significant at the P_adj_=0.05 or (further restricted) P_adj_=0.01 level as well as on the Wald statistic provided by DESeq2 instead of Log2-fold changes. However, we found no improved or worse performance in prediction accuracy when using the Wald statistic or censored sets of genotypes. As highlighted above, this suggests that there is valuable latent information in genotypes whose change in abundance does not meet traditional significance cut-offs. We also trained additional models, where higher-level features (GC content, predicted minimum free energy, base-pairing status at particular rungs of the helix, etc.) were explicitly included. Inclusion did not improve predictive performance, suggesting that emergent properties are captured by models based solely on nucleotide identity at the eight sites. We found that, while inclusion of higher-order features is tempting to increase interpretability, this is a double-edged sword: although higher-order features with large gains can help with interpretation, continuous features or features with more categories can in principle provide more explanatory power for a continuous outcome variable than binary features or features with few categories. Consequently, these features may end up “hogging” predictive power, without necessarily providing greater insight. Exclusive use of nucleotide identities at a given site has the advantage of allowing direct comparison of explanatory power between all features in the model.

To calculate the predictive power of the model (prediction accuracy), one would ordinarily predict fold-change values for the test set (the genotypes left out during training of the model) and compare this to the observed changes. When we do so we obtain correlation coefficients ρ>0.83 for both 30°C and 37°C data. Note that, in terms of the variance of fitness across genotypes explained by the model, this estimate arguably better approximates the genetic variance (V_g_) rather than total phenotypic variance (V_p_=V_g_+V_e_). This is because computing fold-changes across several replicates should reduce the environmental part of the variance (V_e_). To be more conservative, we also calculated fold-changes from five of the six replicates, trained the model on those fold-changes and then tested model performance on the nominal fold-change of the remaining replicate. As expected – given that a single-replicate estimate is bound to be noisier than cross-replicate estimates, correlation coefficients here drop slightly, to ρ>0.63 for both 30°C and 37°C.

### Molecular dynamics simulations

The starting structure for simulations was constructed by templating the sequence of the P1/P1ex region of Tet-119(C20A) onto a previously solved P1/P1ex NMR structure (PDB 1HLX) (Allain and Varani 1995). We then constructed 16 models comprising every single and double base mutation at nucleotides N_2_ and N_21_. All models were parameterized using the Amber RNA OL3 potentials for RNA (Banáš *et al.* 2010), solvated with 14 Å of TIP3P water and neutralized with NaCl. Energy minimization was performed for 2000 steps using combined steepest descent and conjugate gradient methods. Following minimization, 20 ps of classical molecular dynamics (cMD) was performed in the NVT ensemble using a Langevin thermostat (Davidchack *et al.* 2009) to regulate the temperature as we heated up from 0 to 300 K. Following the heat-up phase, we preformed 100 ns of cMD in the isobaric/isothermal (NPT) ensemble using the Berendsen barostat (Berendsen *et al.* 1998) to maintain constant pressure during the simulation. All simulations were preformed using GPU (CUDA) Version 18.0.0 of PMEMD (Götz *et al.* 2012; Le Grand *et al.* 2013; Salomon-Ferrer *et al.* 2013) with long-range electrostatic forces treated with Particle-Mesh Ewald summation. RNA base pair properties were calculated using CPPTRAJ (Roe and Cheatham 2013) and visualized using VMD (Humphrey *et al.* 1996).

## ACKNOWLEDGEMENTS

We thank Feng Guo (UCLA) for the gift of Tet-199(C20A) and mutant plasmids, the LMS Genomics facility for library construction and sequencing, members of the Molecular Systems lab and Romain Strock for discussion, and Ben Lehner, Peter Sarkies, and Karen Sarkisyan for comments on the manuscript. This work was supported by Medical Research Council core funding to TW, a Marie Skłodowska-Curie grant (agreement no. 747199) to VWCS, and a UKRI Innovation Fellowship to JBS. This project made use of time on UK Tier 2 JADE granted via the UK High-End Computing Consortium for Biomolecular Simulation, HECBioSim (http://hecbiosim.ac.uk), supported by EPSRC (grant no. EP/R029407/1).

---

**Figure S1.**
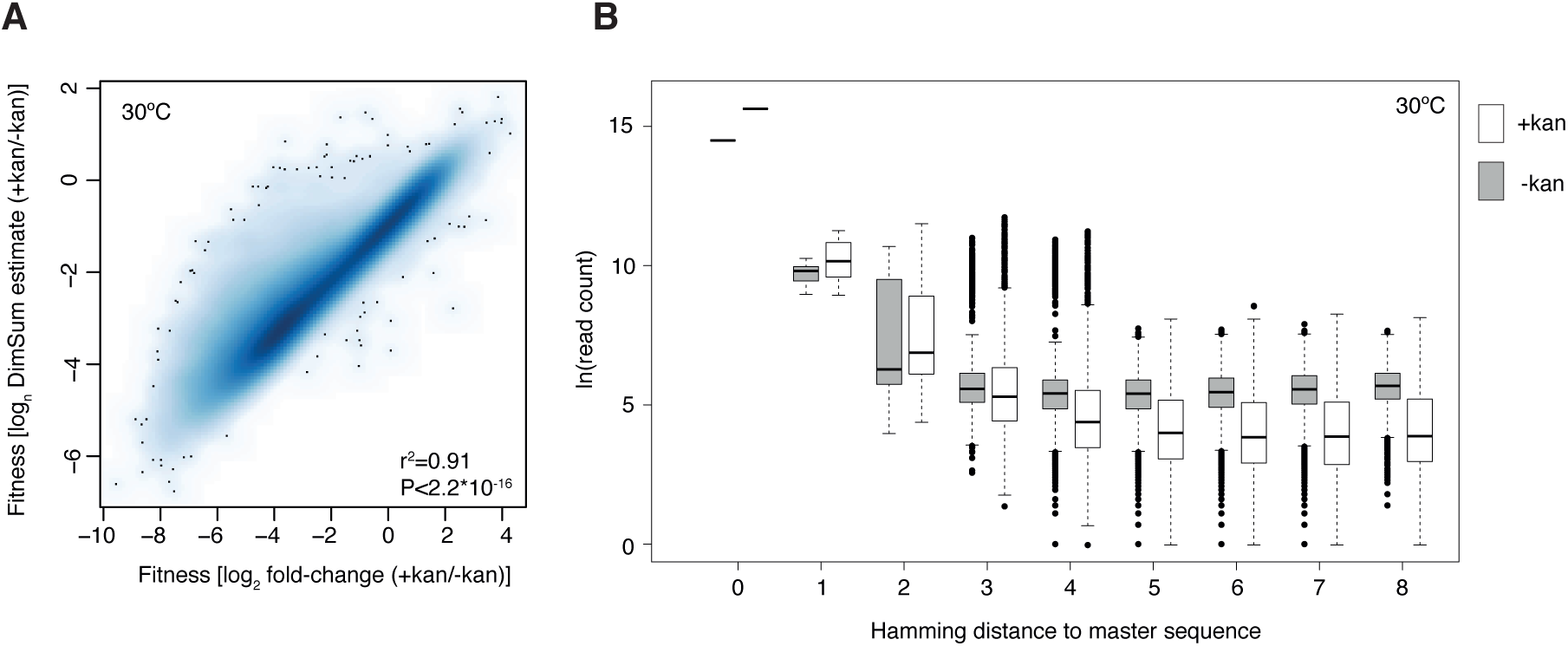
(**A**) Correlation of fitness estimates derived from the DiMSum pipeline and using the DESeq2 framework. (**B**) Biased distribution of read counts prior to and after selection. As a consequence of library generation, genotypes closer to the master sequence are, on average, more common even prior to selection.

**Figure S2.**
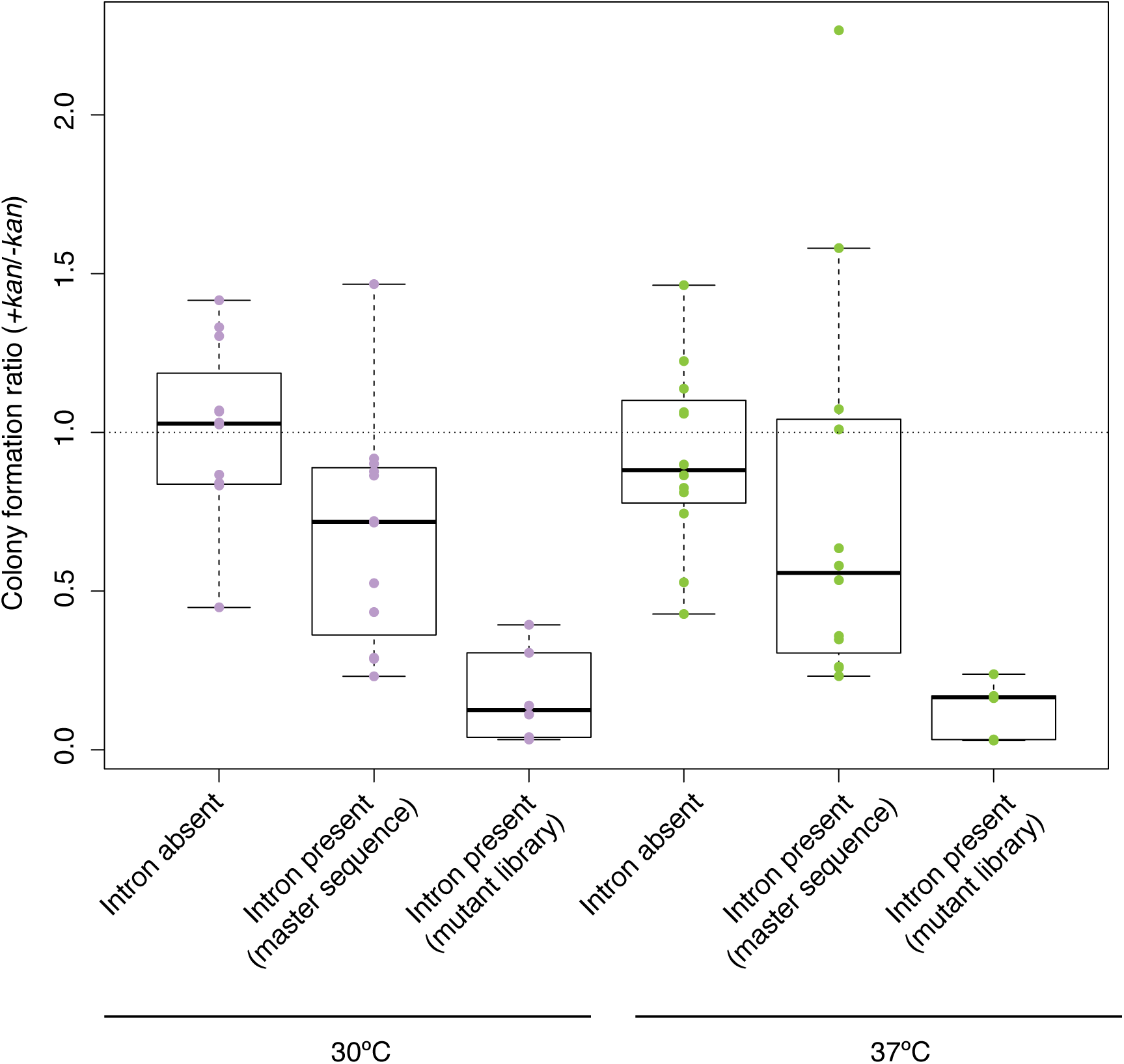
The effect of intron insertion into *knt* on colony formation in *E. coli.*

**Figure S3.**
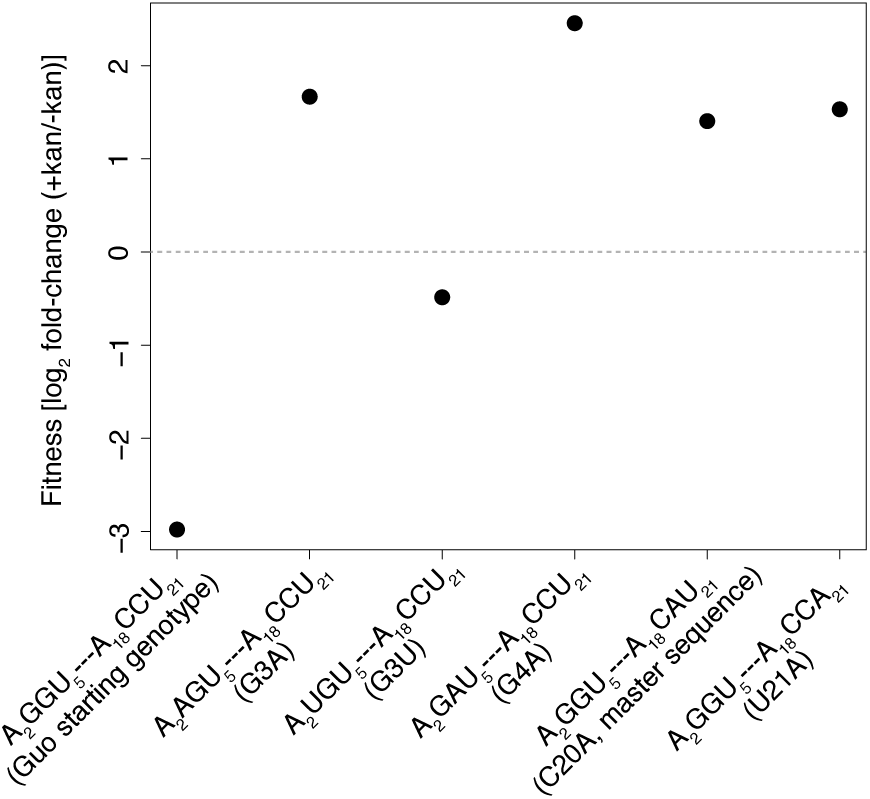
Relative fitness of the Tet-119 genotype and previously described single-mutation derivatives, including our master sequence Tet-119(C20A) (Guo and Cech 2002). All derivatives have previously been shown to have higher splicing activity than Tet-119 and all exhibit higher fitness in our assay.

**Figure S4.**
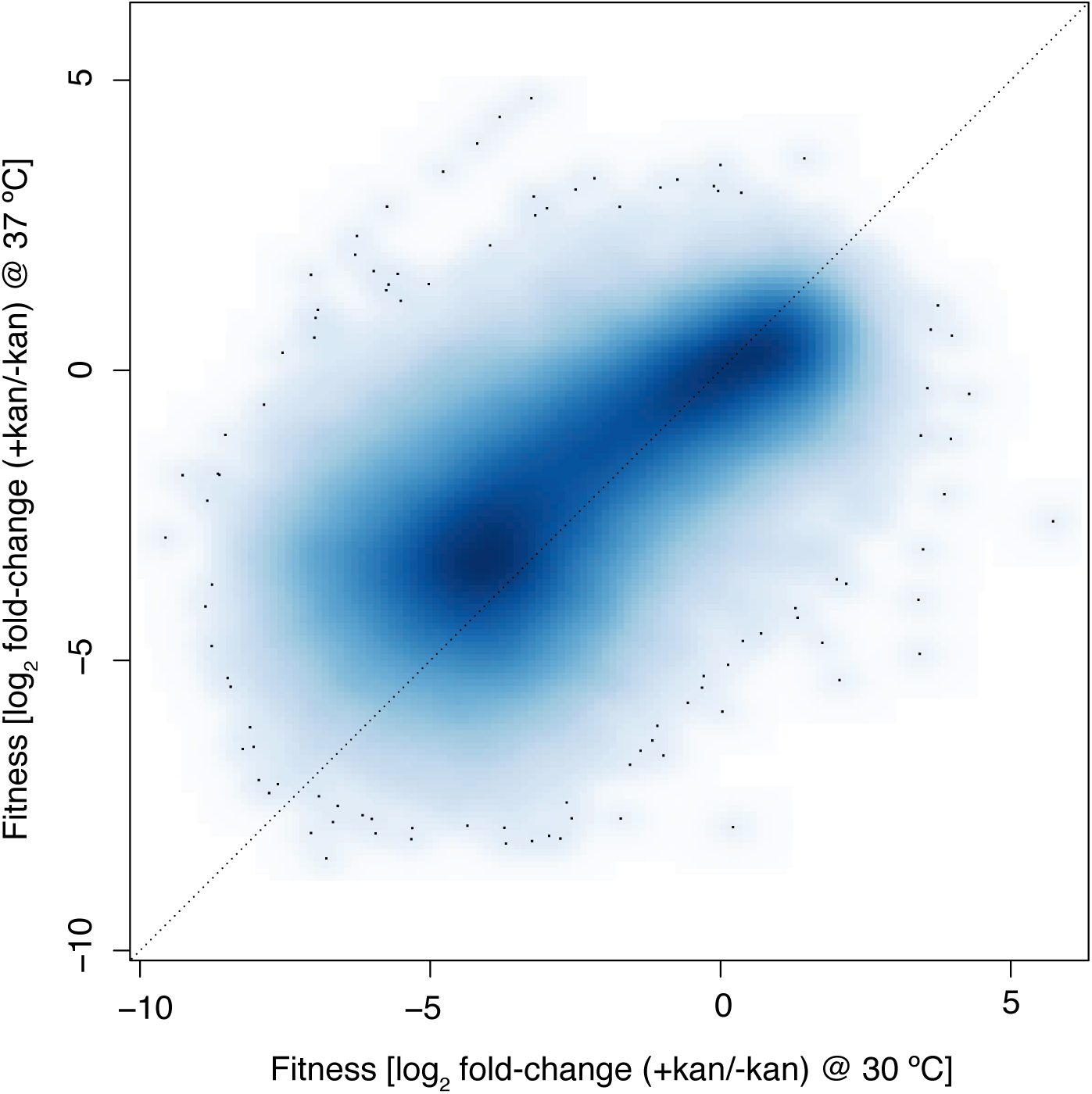
Correlation of fitness effects at 37°C and 30°C (ϱ=0.75, P<2.2*10^−16^).

**Figure S5.**
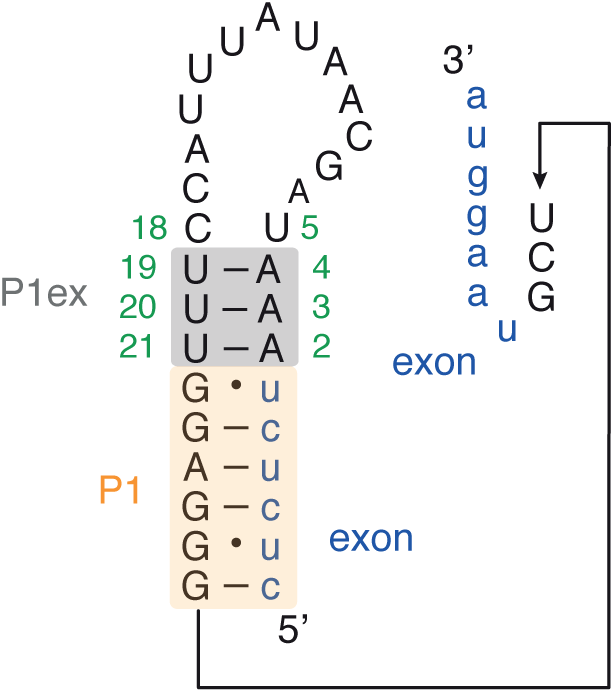
The sequence and secondary structure of P1 and P1ex in the native *Tetrahymena thermophila* group I intron and its pre-rRNA environment.

**Figure S6.**
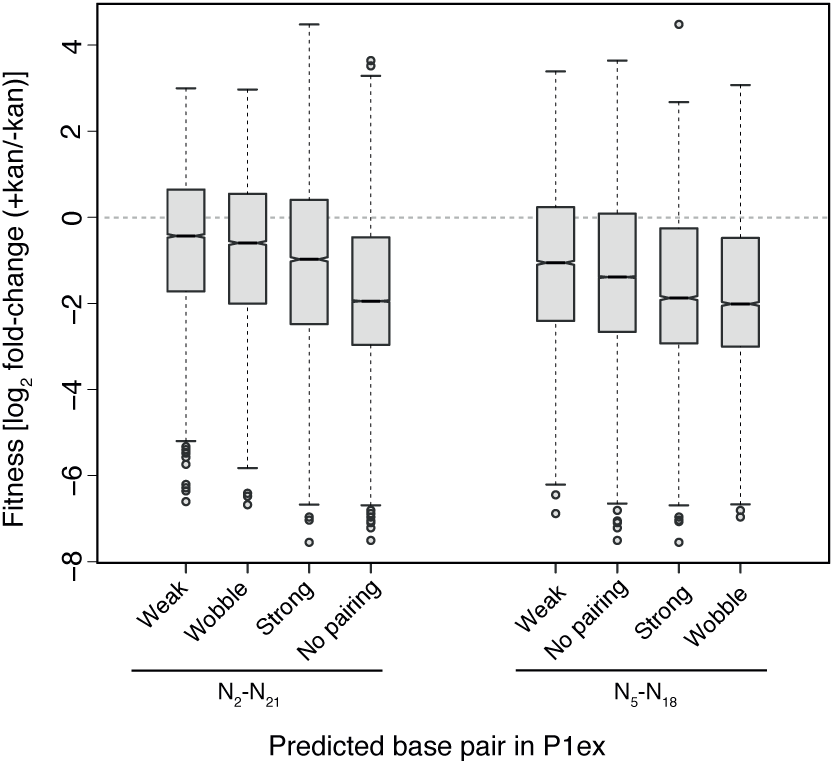
Fitness binned according to the types of on-target base-pairing interactions that can be formed by N_2_-N_21_ and N_5_-N_18_.

**Figure S7.**
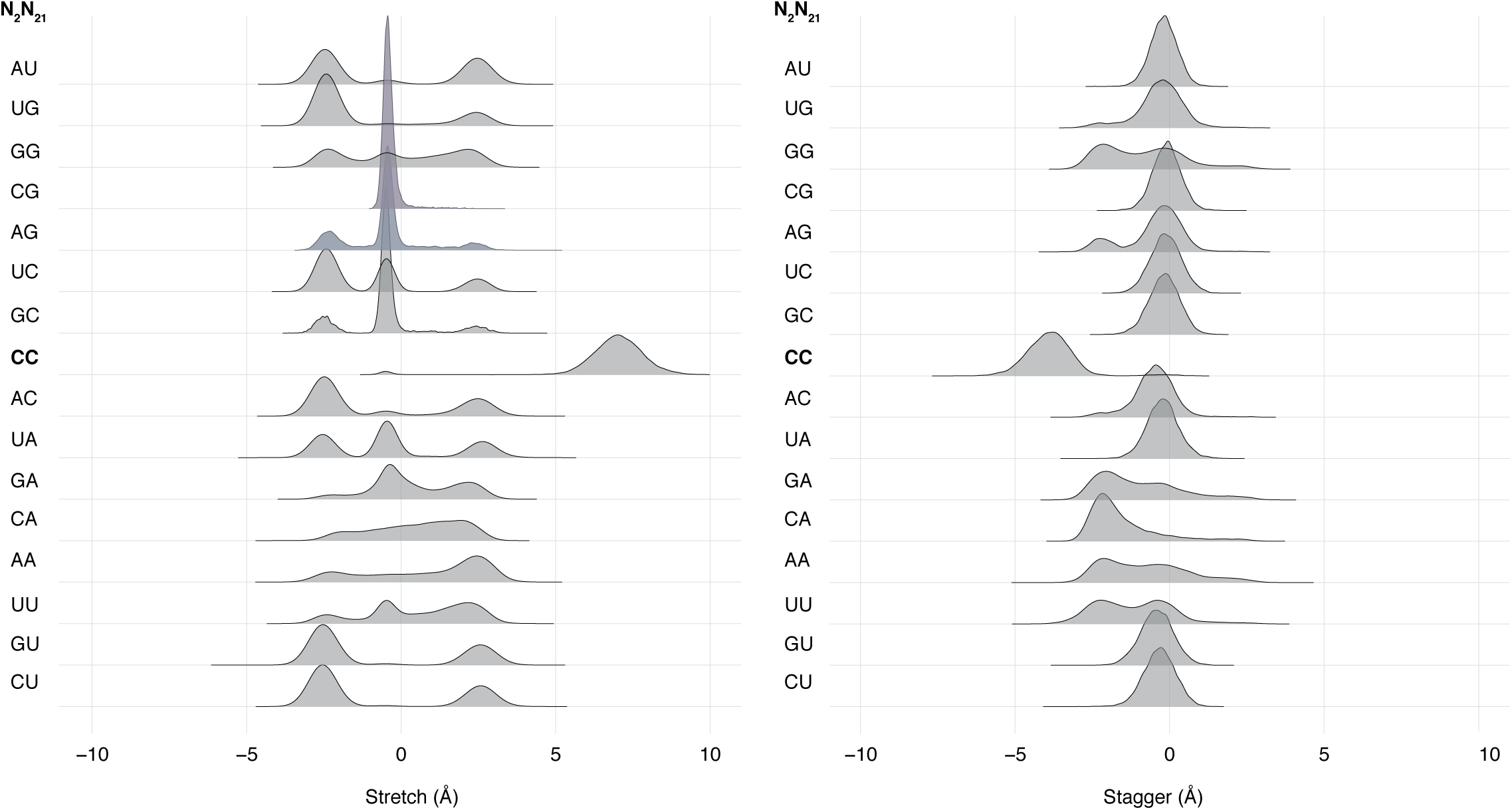
Stretch and stagger measured at the splice site (U_1_-G_22_) for all possible nucleotide combinations at N_2_/N_21_.

**Figure S8.**
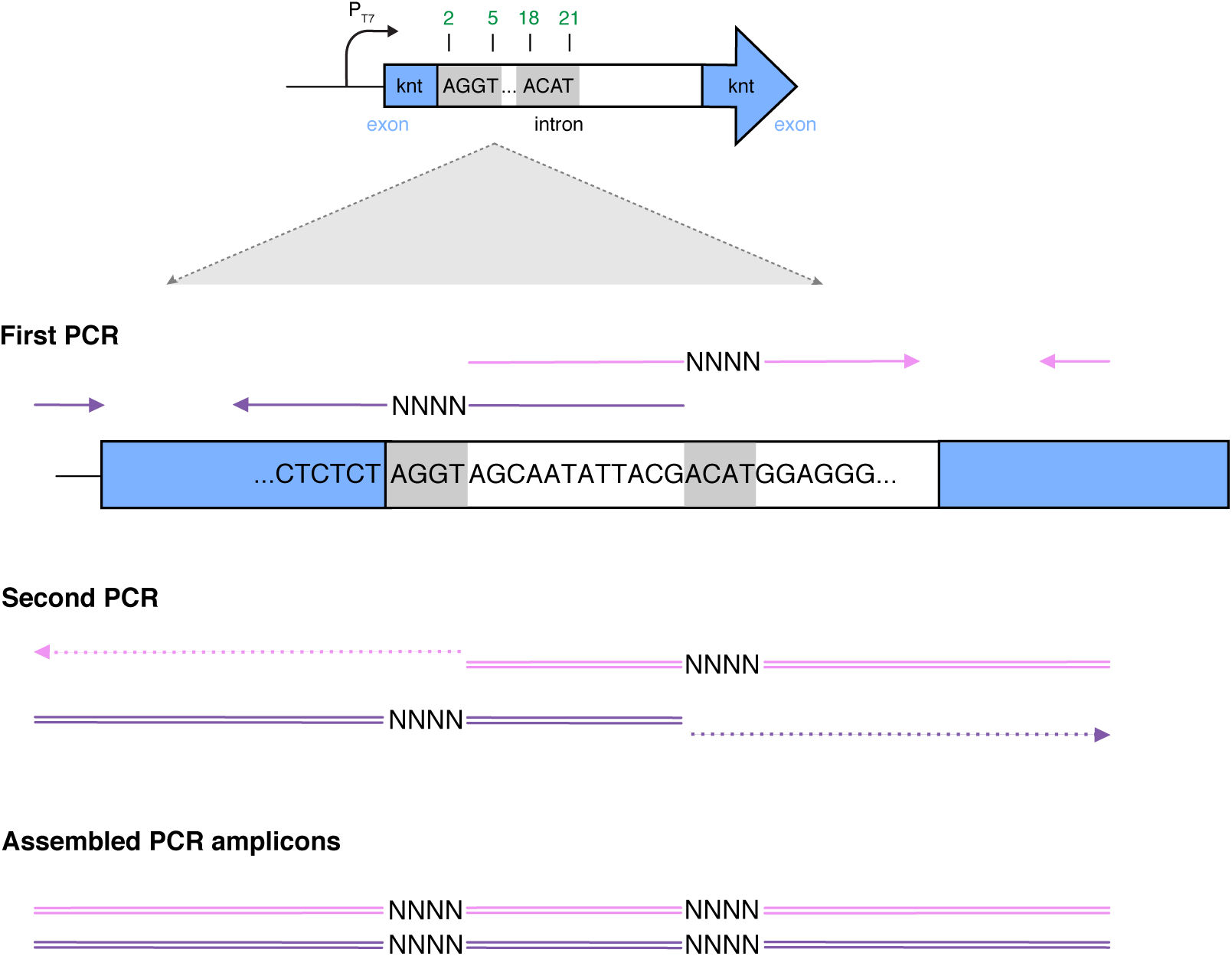
Generation of mutant library using site-saturation mutagenesis via a two-step PCR. Solid arrows denote oligonucleotides. In the first step, two pairs of oligonucleotides containing mixed bases (https://www.idtdna.com/pages/products/custom-dna-rna/mixed-bases) are used to amplify two separate fragments (purple and pink) containing one P1ex sub-region each. The 3’ end of the purple fragment and the 5’ end of the pink fragment share a 12-bp overlapping region, which allow self-annealing and subsequent 3’ extension during the second PCR. As a result, the assembled amplicons contain two varying sub-regions. The purple fragment is amplified using primers Frag1-f and Frag1-r, whereas the pink fragment is amplified using primers Frag2-f and Frag2-r (Table S3).

**Table S1. Fitness of individual genotypes across conditions.** (see supplementary file)

**Table S2.**
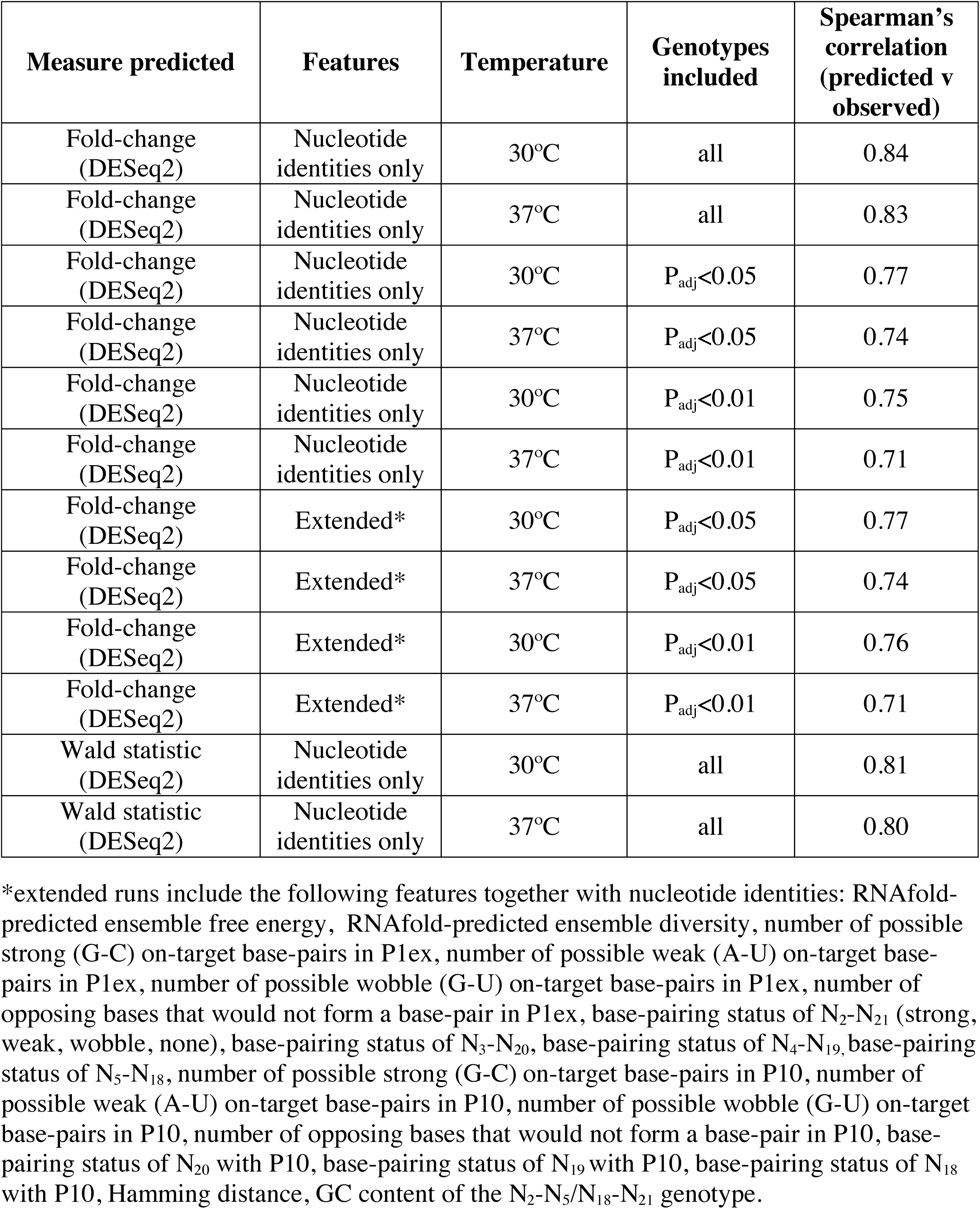
XGBoost models.

**Table S3.**
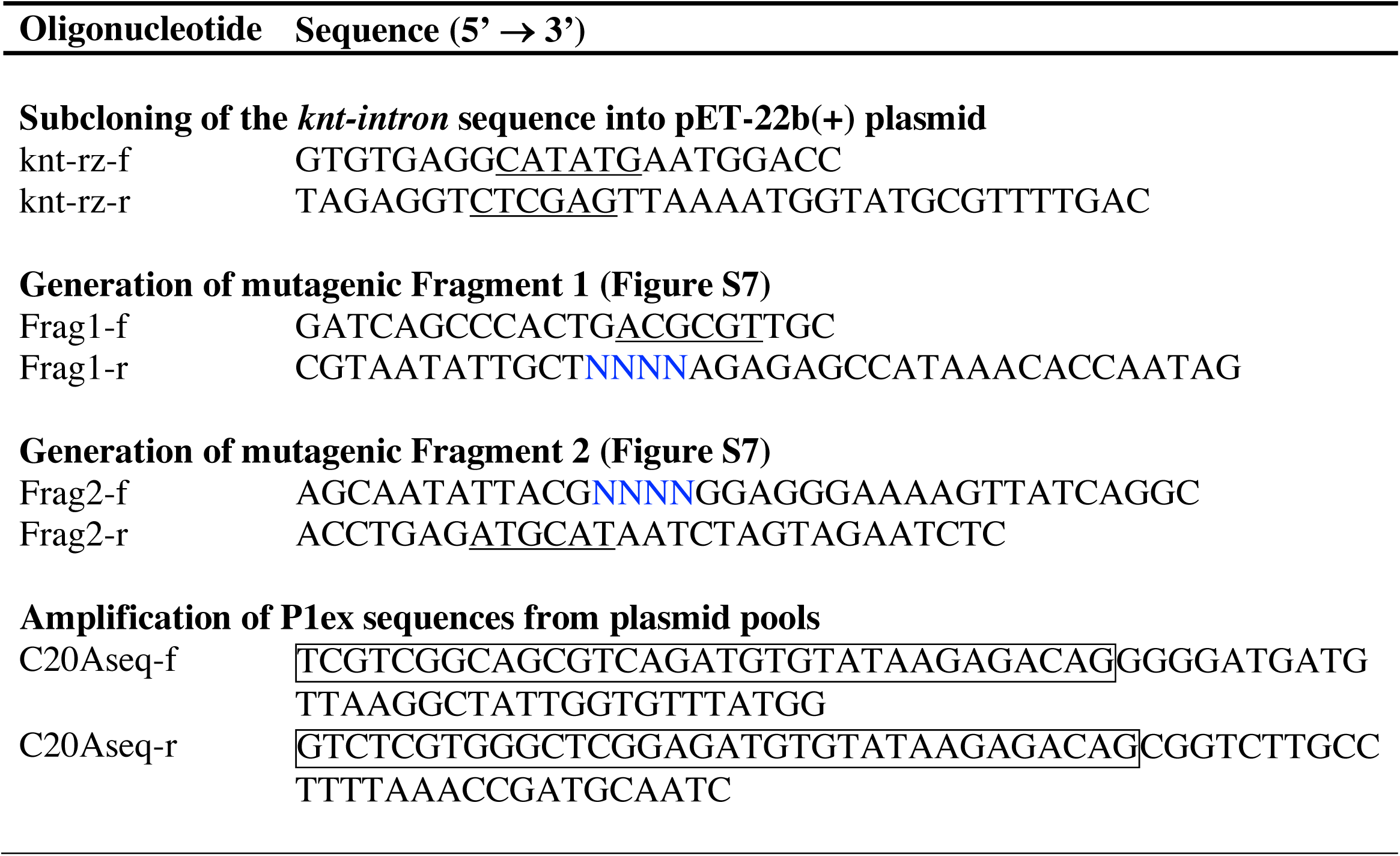
Oligonucleotides used in this study. “f” indicates a forward primer and “r” indicates a reverse primer. Mixed bases are in blue. The restriction sites are underlined. The Illumina adapter sequences are boxed.

**File S1.** Molecular dynamics simulation of C_2_/C_21_ genotype, highlighting rotation of U_1_ out of the helix core.

## Notes

### Competing Interest Statement

The authors have declared no competing interest.

